# Multi-Modality Machine Learning Predicting Parkinson’s Disease

**DOI:** 10.1101/2021.03.05.434104

**Authors:** Mary B. Makarious, Hampton L. Leonard, Dan Vitale, Hirotaka Iwaki, Lana Sargent, Anant Dadu, Ivo Violich, Elizabeth Hutchins, David Saffo, Sara Bandres-Ciga, Jonggeol Jeff Kim, Yeajin Song, Matt Bookman, Willy Nojopranoto, Roy H. Campbell, Sayed Hadi Hashemi, Juan A. Botia, John F. Carter, Melina Maleknia, David W. Craig, Kendall Van Keuren-Jensen, Huw R. Morris, John A. Hardy, Cornelis Blauwendraat, Andrew B. Singleton, Faraz Faghri, Mike A. Nalls on behalf of the Accelerating Medicines Program - Parkinson’s Disease (AMP PD) and the Global Parkinson’s Genetics Program (GP2).

## Abstract

**Background:** Personalized medicine promises individualized disease prediction and treatment. The convergence of machine learning (ML) and available multi-modal data is key moving forward. We build upon previous work to deliver multi-modal predictions of Parkinson’s Disease (PD).

**Methods:** We performed automated ML on multi-modal data from the Parkinson’s Progression Marker Initiative (PPMI). After selecting the best performing algorithm, all PPMI data was used to tune the selected model. The model was validated in the Parkinson’s Disease Biomarker Program (PDBP) dataset. Finally, networks were built to identify gene communities specific to PD.

**Findings:** Our initial model showed an area under the curve (AUC) of 89.72% for the diagnosis of PD. The tuned model was then tested for validation on external data (PDBP, AUC 85.03%). Optimizing thresholds for classification, increased the diagnosis prediction accuracy (balanced accuracy) and other metrics. Combining data modalities outperforms the single biomarker paradigm. UPSIT was the largest contributing predictor for the classification of PD. The transcriptomic data was used to construct a network of disease-relevant transcripts.

**Interpretation:** We have built a model using an automated ML pipeline to make improved multi-omic predictions of PD. The model developed improves disease risk prediction, a critical step for better assessment of PD risk. We constructed gene expression networks for the next generation of genomics-derived interventions. Our automated ML approach allows complex predictive models to be reproducible and accessible to the community.

**Funding:** National Institute on Aging, National Institute of Neurological Disorders and Stroke, the Michael J. Fox Foundation, and the Global Parkinson’s Genetics Program.

**RESEARCH IN CONTEXT:** *Evidence before this study:* Prior research into predictors of Parkinson’s disease (PD) has either used basic statistical methods to make predictions across data modalities, or they have focused on a single data type or biomarker model. We have done this using an open-source automated machine learning (ML) framework on extensive multi-modal data, which we believe yields robust and reproducible results. We consider this the first true multi-modality ML study of PD risk classification.

*Added value of this study:* We used a variety of linear, non-linear, kernel, neural networks, and ensemble ML algorithms to generate an accurate classification of both cases and controls in independent datasets using data that is not involved in PD diagnosis itself at study recruitment. The model built in this paper significantly improves upon our previous models that used the entire training dataset in previous work^1^. Building on this earlier work, we showed that the PD diagnosis can be refined using improved algorithmic classification tools that may yield potential biological insights. We have taken careful consideration to develop and validate this model using public controlled-access datasets and an open-source ML framework to allow for reproducible and transparent results.

*Implications of all available evidence:* Training, validating, and tuning a diagnostic algorithm for PD will allow us to augment clinical diagnoses or risk assessments with less need for complex and expensive exams. Going forward, these models can be built on remote or asynchronously collected data which may be important in a growing telemedicine paradigm. More refined diagnostics will also increase clinical trial efficiency by potentially refining phenotyping and predicting onset, allowing providers to identify potential cases earlier. Early detection could lead to improved treatment response and higher efficacy. Finally, as part of our workflow, we built new networks representing communities of genes correlated in PD cases in a hypothesis-free manner, showing how new and existing genes may be connected and highlighting therapeutic opportunities.

## INTRODUCTION

For progressive neurodegenerative diseases, early and accurate diagnosis is key to effectively developing and using new interventions. This early detection paradigm aims to identify, analyze, and prevent or manage the disease before the patient recognizes signs and symptoms while the disease process is most amenable to intervention. Here we describe work that facilitates accurate and early diagnosis using cost-effective methods in a data-driven manner ^1^.

Biomedical researchers are currently at the convergence of scientific advances that will facilitate progress in early detection and remote identification of potentially high-risk individuals: first, the availability of substantial clinical, demographic, and genetic/genomic datasets. Second, advances in the automation of machine learning (ML) pipelines and artificial intelligence, to maximize the value of this massive amount of readily available data ^2^. Previous biomarker studies, particularly in neurodegenerative diseases, have focused on widely known statistical approaches and linear models, using a single metric or handful of metrics for predictions. We aim to contribute to the field with non-linear and ML-based approaches and leverage rapidly growing publicly available data to build these models. We also extend these models, providing not just disease prediction but also biological insight.

We have used publicly available multi-modal Parkinson’s disease (PD) data to build an accurate peri-diagnostic model to predict disease risk. We also used the features nominated by our workflow to build novel, unbiased networks of genes related to the onset of PD that highlight biological pathways of interest and therapeutic targets. This work leveraged clinico-demographic and multi-omic data produced and curated to build models that can impact both trial recruitment and drug development. The models we have developed here improved performance over previous related efforts, with performance metrics at current cross-validation in withheld samples being equivalent, or in some cases, better than the training phase of earlier work^1^. The data came from the Accelerating Medicines Program - Parkinson’s Disease (AMP PD) program [https://amp-pd.org/] and the code used to carry out analyses comes from open-source automated ML software, all of which have been made publicly available to support reproducibility, transparency, and open science ^3,4^.

## METHODS

### Online Appendix

Full methods and links to code can be found in the **Online Appendix**. A brief overview of the process can be found below. All of the data and code used for this project is also available at publication via a linked Terra workspace for full reproducibility [link pending publication]. GenoML is an open-source Python package automating machine learning workflows for genomics^5^. Source code and documentation is available at [https://genoml.com/] and on GitHub [https://github.com/GenoML/genoml2]. AMP PD data and quality control notebooks are access-controlled [https://amp-pd.org/]. Additionally, we have developed an interactive website [https://share.streamlit.io/anant-dadu/shapleypdpredictiongenetics/main] where researchers can investigate components of the predictive model.

### Procedures and statistical analysis overview

Figure 1. summarizes the workflow and data used in this project. Our workflow began with data munging that includes feature selection, adjustment, and normalization. Then we moved on to algorithm competition and feature selection based on a 70:30 (training:testing) split in the PPMI dataset. We then compared how each algorithm performed on identical training and testing data. Once the best performing algorithm had been selected, a thorough hyperparameter tuning of the algorithm with 5 fold cross-validation (also in the entire PPMI) was performed. Model was exported to enable external validation and transfer learning in the readers’ own data. This hyperparameter tuning and cross-validation phase was carried out to both improve performance but also reduce bias ^6^. We validated the models built by taking the trained and tuned models from PPMI and fit them to the external validation dataset, PDBP. Cohort summaries can be found below in **Table 1** with additional information in the **Online Appendix**.

**Figure 1:**
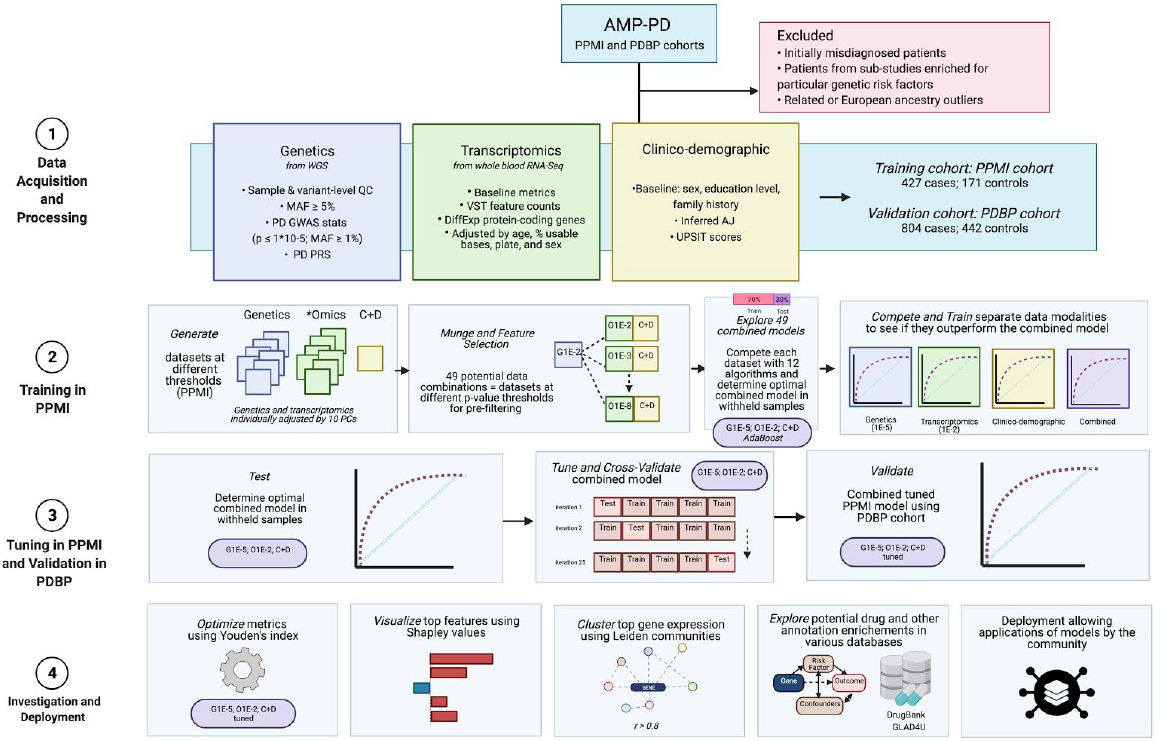
Workflow and Data Summary. Scientific notation in the workflow diagram denotes minimum p-values from reference GWAS or differential expression studies as a pre-screen for feature inclusion.

**Table 1:** Descriptive statistics of studies included from AMP PD.

| **Study** | **Status** | **Age at baseline mean (SD)** | **UPSIT score mean (SD)** | **Male (%)** | **Positive family history of PD (%)** | **Inferred Ashkenazi ancestry (%)** |
| --- | --- | --- | --- | --- | --- | --- |
| *PPMI* | *Case* | 61.75 (9.69) | 23.48 (8.35) | 65.57 | 25.53 | 6.09 |
|  | *Control* | 60.61 (10.43) | 34.18 (4.71) | 63.74 | 5.85 | 11.11 |
| *PDBP* | *Case* | 64.59 (8.99) | 19.65 (8.01) | 64.18 | 24.88 | 3.61 |
|  | *Control* | 62.87 (10.96) | 32.52 (5.98) | 45.25 | 8.14 | 4.07 |

### Role of the funding source

The study’s funders had no role in the study design, data collection, data analysis, data interpretation, or writing of the report. All authors and the public can access all data and statistical programming code used in this project for the analyses and results generation. MAN takes final responsibility for the decision to submit the paper for publication.

## RESULTS

We have shown that integrating multiple modalities improved model performance in predicting PD diagnosis in a mixed population of cases and controls. Additional interpretation for ML metrics and models is included in the **Online Appendix**. Our multi-modality model showed a higher area under the curve (AUC; 89.72%) than just the clinico-demographic data available prior to neurological assessment (87.52%), the genetics-only model from genome sequencing data and polygenic risk score (PRS; 70.66%), or the transcriptomics-only model from genome-wide whole blood RNA sequencing (RNAseq) data (79.73%) in withheld PPMI samples (see **Table 2** and **Figure 2** for summaries).This model’s performance improved in both PPMI and PDBP after tuning, described below and in **Table 3**. Similar results can be seen when this model is validated in the PDBP data set (AUC from the combined modality model at 83.84% before tuning) detailed in **Table 4** and **Figure 3**. Additionally, the multi-modal model also had the lowest false positive and false negative rates compared to other models, only focusing on a single modality, in both the withheld test set in PPMI and in the PDBP validation set. Thus, moving from single to multiple data modalities yielded better results in not only AUC but across all performance metrics. A strength of using multi-modal approaches is that some modalities may better predict case or control status than others (**Table 2**). Here, we leveraged data diversity to increase overall sensitivity and specificity. Our final multi-modal model had higher accuracy and balanced accuracy at 85.56% and 82.41%, respectively, sensitivity at 89.31%, and specificity at 75.51%, when compared to models built only on a single data modality. Notably, this improved balanced accuracy is of particular importance in binary classifiers where one of the predicted classes is much rarer than the other, like PD, which is relatively infrequent in the general population. Special attention was given to validate the model, interpreting and visualizing the top features aiding in the prediction of classification, and further investigation into optimizing the model, developing hypothesis-free transcriptomic communities, and exploring potential drug-gene interactions.

**Table 2:** Performance metric summaries comparing training in withheld samples in PPMI.

| Data Modality | Stage | Algorithm | AUC (%) | Accuracy (%) | Balanced accuracy (%) | Log Loss | Sensitivity | Specificity | PPV | NPV |
| --- | --- | --- | --- | --- | --- | --- | --- | --- | --- | --- |
| Genetics (P<1E-5) | Training in PPMI (70:30) | MLPClassifier | 70.66 | 70.00 | 60.64 | 0.83 | 0.83 | 0.38 | 0.77 | 0.48 |
| Clinico-demographic | Training in PPMI (70:30) | LogisticRegression | 87.52 | 79.44 | 75.27 | 0.39 | 0.85 | 0.65 | 0.86 | 0.64 |
| Transcriptomics (P<1E-2) | Training in PPMI (70:30) | SVC | 79.73 | 73.89 | 54.60 | 0.48 | 0.97 | 0.12 | 0.75 | 0.60 |
| Combined | Training in PPMI (70:30) | AdaBoostClassifier | 89.72 | 85.56 | 82.41 | 0.63 | 0.89 | 0.76 | 0.91 | 0.73 |

**Figure 2:**
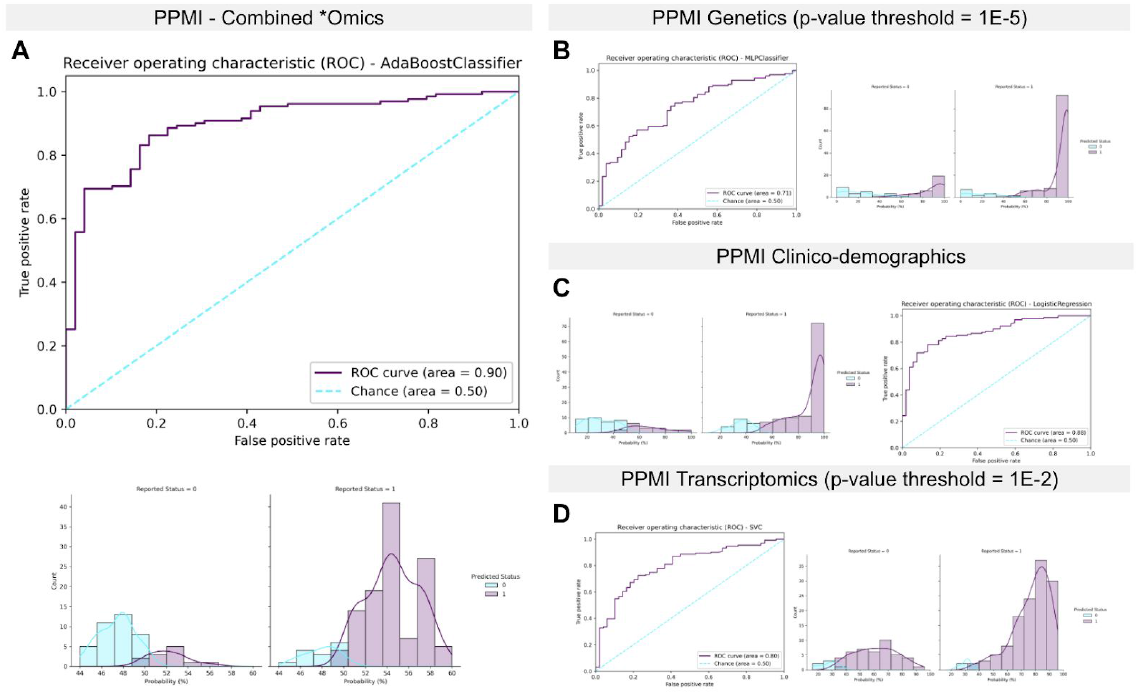
Receiver operating characteristic curves and case probability density plots in withheld training samples at default thresholds comparing performance metrics in different data modalities from the PPMI dataset. P-values mentioned indicate the threshold of significance used per data type, except for the inclusion of all clinico-demographic features.

**Table 3:** Performance metric summaries comparing tuning in PPMI for different subsets of data modalities.

| Data Modality | Stage | Algorithm | AUC at training (%) | Mean, AUC during CV for baseline model (%) | Standard deviation, AUC during CV for baseline model (%) | Min, AUC during CV for baseline model (%) | Max, AUC during CV for baseline model (%) | Mean, AUC during CV for tuned model (%) | Standard deviation, AUC during CV for tuned model (%) | Min, AUC during CV for tuned model (%) | Max, AUC during CV for tuned model (%) |
| --- | --- | --- | --- | --- | --- | --- | --- | --- | --- | --- | --- |
| Genetics (P<1E-5) | Tuning in PPMI | MLPClassifier | 70.66 | 69.44 | 4.46 | 62.45 | 75.73 | 70.93 | 5.39 | 61.29 | 76.71 |
| Clinico-demographic | Tuning in PPMI | LogisticRegression | 87.52 | 88.51 | 2.17 | 86.19 | 91.98 | 88.55 | 2.20 | 86.33 | 92.15 |
| Transcriptomics (P<1E-2) | Tuning in PPMI | SVC | 79.73 | 78.05 | 4.27 | 71.49 | 82.62 | 79.01 | 4.71 | 70.88 | 84.01 |
| Combined | Tuning in PPMI | AdaBoostClassifier | 89.72 | 86.99 | 2.30 | 84.27 | 90.70 | 90.17 | 1.64 | 88.06 | 92.73 |

**Table 4:** Performance metric summaries comparing combined tuned and untuned model performance on the PDBP validation dataset.

| Data Modality | Stage | Algorithm | AUC (%) | Accuracy (%) | Balanced Accuracy (%) | Log Loss | Sensitivity | Specificity | PPV | NPV |
| --- | --- | --- | --- | --- | --- | --- | --- | --- | --- | --- |
| Combined | Untuned in PPMI as reference | AdaBoostClassifier | 89.72 | 85.56 | 82.41 | 0.63 | 0.89 | 0.76 | 0.91 | 0.73 |
| Combined; Untuned | Validation in PDBP | AdaBoostClassifier | 83.84 | 75.81 | 69.31 | 0.64 | 0.93 | 0.46 | 0.75 | 0.78 |
| Combined; Tuned | Validation in PDBP | AdaBoostClassifier | 85.03 | 75.00 | 68.09 | 0.67 | 0.93 | 0.43 | 0.74 | 0.78 |

**Figure 3:**
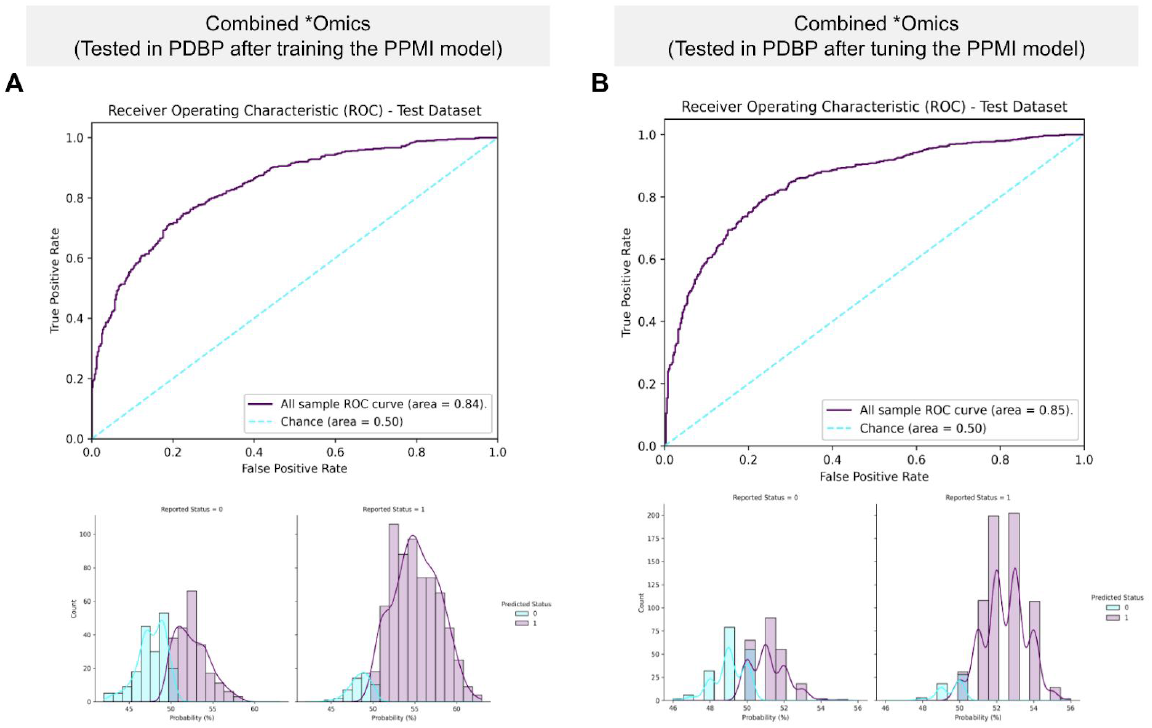
Receiver operating characteristic and case probability density plots in the external dataset (PDBP) at validation for the trained and then tuned models at default thresholds. Probabilities are predicted case status (r1), so controls (a status of 0) skews towards more samples on the left, and positive PD cases (a status of 1) skews more samples on the right.

One benefit of the ML approach we have used is its ability to tune model parameters. The best performing tuned model that included all data showed an AUC distribution of 88.06% to 92.70% at 5-fold cross-validation with a mean of 90.20% and a standard deviation of 2.3% in PPMI (see **Table 3**). When validated in the PDBP data, we saw an AUC of 85.03%, sensitivity at 93.12%, and specificity at 43.07% for the tuned multi-modal model.

These models then improved further when *post-hoc* optimization of case probability thresholds was carried out. We considered the optimized version of the tuned model (including all data modalities) as our gold standard. When applied to withheld PPMI samples, the training phase model increased its balanced accuracy quantified performance to 83.95%. This optimization also led to improved balanced accuracy of 77.97% when fitting the tuned model referenced above to the PDBP validation data. See **Table 5** for details on other related metrics and a summary of optimized versus default thresholds. In general, our threshold optimization allowed general increases in classifier performance at a minimal computational cost.

**Table 5:** Optimizing the AUC threshold in withheld training samples and in the validation data.

| Dataset | Model | Optimization | Case Probability Threshold (%) | Accuracy (%) | Balanced accuracy (%) | Log loss | Sensitivity | Specificity | PPV | NPV |
| --- | --- | --- | --- | --- | --- | --- | --- | --- | --- | --- |
| PPMI, withheld samples | Training phase | optimized | 51 | 85 | 83.95 | 0.05 | 0.86 | 0.82 | 0.93 | 0.69 |
| PPMI, withheld samples | Training phase | default | 50 | 85.56 | 82.41 | 0.05 | 0.89 | 0.76 | 0.91 | 0.73 |
| PDBP, external test samples | Tuned model | optimized | 51 | 78.58 | 77.97 | 0.07 | 0.80 | 0.76 | 0.85 | 0.68 |
| PDBP, external test samples | Tuned model | default | 50 | 75 | 68.09 | 0.09 | 0.93 | 0.43 | 0.74 | 0.78 |

Our model build included 71 SNPs and 596 protein-coding transcripts in addition to expected features like the demographics, family history, olfactory function, and previous genome-wide significant polygenic risk estimates in the form of PRS^7^. The Shapley Additive exPlanation (SHAP) plots in **Figure 4** show the relative importance of the features in the model approximated using withheld training data. When investigating the SHAP values for both the training and testing samples, the UPSIT score, as well as PRS, contributed most to the predictive power of the model, but the accuracy of these are supplemented by many smaller effect transcripts and risk SNPs. It also indicates that the lower UPSIT score (designated by the blue color on the left-hand side) value corresponds to a higher probability of PD, as most of the blue-colored points lie on the right side of the baseline risk estimate. Looking closer at these features, we can also observe that the directionality of different genetic features is not uniform. This signifies that over-expression of some genes corresponds to healthy controls while for some features it is in the opposite direction. We have also created a website that allows readers to further explore feature contributions to model accuracy in various scenarios [https://share.streamlit.io/anant-dadu/shapleypdpredictiongenetics/main]. The addition of SNPs outside the PRS could suggest potential compensatory or risk modification effects interacting with the PRS. For more information on pairwise interactions between the PRS and individual SNPs, please see the **Online Appendix**.

**Figure 4:**
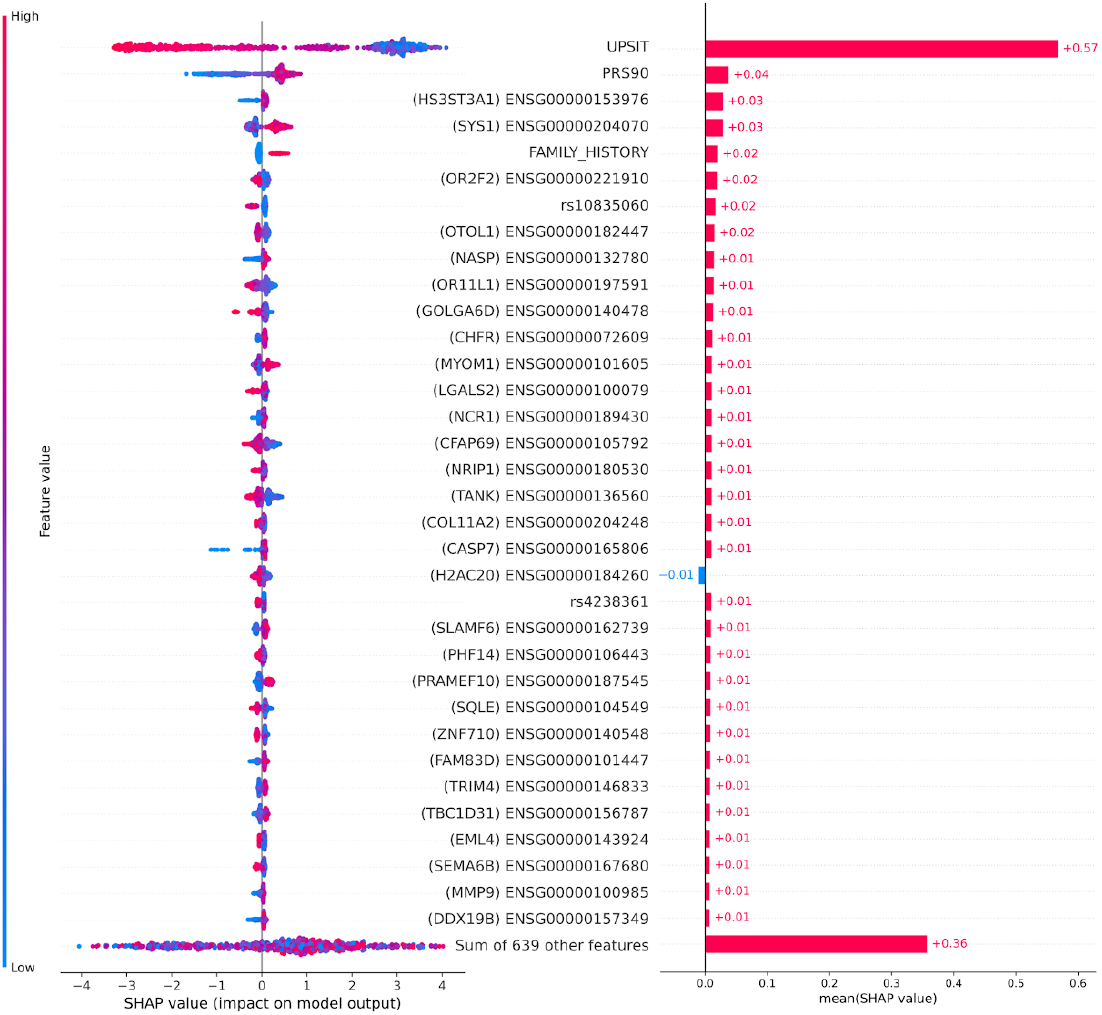
Feature importance plots for top 5% of features in withheld training data.

Gene expression network communities were constructed using RNAseq data extracted from positive PD cases. These genes were nominated by the feature selection process. These communities of genes represent part of a potentially novel PD-specific network derived from whole blood RNAseq. Consider the network itself to be conceptually similar to a pathway, composed of genes whose expression was strongly correlated in the case-only transcriptomics dataset, and the communities being subgroups of closely related genes within the larger set. We identified 13 network communities consisting of 300 genes with an Erdos-Renyi modularity score of 0.794 (a modularity score closer to 1 indicates better model fit). A link to the full community annotations and a graphical summary can be found in the **Online Appendix**.

For genes in our network communities, we evaluated the potential over representation of known drug target genes across the identified communities. When comparing the genes defined as part of the network communities (N = 300) to those selected for inclusion as features in the case:control model build (N = 598), we noted enrichments of genes connected to fostamatinib (FDR adjusted p-value 2.21e-4, for genes *MYLK, EPHA8, HCK, DYRK1B*, and *BUB1B-PAK6*) and copper (FDR adjusted p-value 0.0286, for genes *HSP90AA, CBX5* and *HSPD1*) from the DrugBank annotations. The same query in the GLAD4U database resulted in a significant over representation of l-lysine annotated genes (FDR adjusted p-value 0.0057, for genes *DDX50, UBA2, ESCO1, CDC34, ANKIB1, PCMT1, DNAJA1, PRMT3, ASPSCR1, BRDT, LOXL4, CBX5, HAT1, MARCH1, HSP90AA1, KPNB1, KMT5B, PSIP1, XPOT, SLC7A9, ZNF131, DDX18, RBBP5*, and *MSL1*). All other variations of our drug target enrichment analyses yielded no significant drugs overrepresented after multiple test correction, although the top ranked result was consistently gamma hydroxybutyric acid (unadjusted p-values 0.0056-0.0001 for genes *SLC16A7, SLC16A3*, and *GABBR1*).

## DISCUSSION

In an era where genetics and genomics combined with clinico-demographic data are increasingly available to researchers, we can now build multi-modal models at a scale that use multiple data modalities for increased performance.

This study illustrates that integrating diverse data modalities into modeling efforts can improve the quality of predictions. This paper is a proof of concept for future directions in this area of predictive modeling in large healthcare data and indicative of its relevance to other areas such as clinical trial enrollment and stratification. In our modeling process, we have succeeded in building robust model(s) of peri-diagnostic PD while also generating *de novo* network communities of genes correlated in PD cases, providing further data for potential therapeutic development. A strength of using this multi-modal approach is that the different modalities compensate for one another, with some modalities better at predicting case status while others are better at classifying controls. All of this was accomplished in a completely transparent and open science framework, from the underlying data to code and resulting models.

While we do not suggest this as a replacement for current diagnostic screening methods, it can be a potential adjunct screening that could aid in identifying high-risk individuals, especially on a large biobank or study recruitment scale. Additional studies will need to be conducted to ascertain the model’s ability to distinguish very early PD cases from other diseases within high risk cohort studies. The estimated prevalence of PD in an aging population is about 2% ^8^. At this prevalence for our optimized PPMI model described in **Table 5**, the positive predictive value (PPV) and negative predictive values (NPV) were calculated to be 8.75% and 99.66%, respectively. The false discovery rate (FDR) and false omission rates (FOR) at this prevalence are 91.26% and 0.34%, respectively. The low FOR indicates that for every 1000 individuals who are classified as healthy controls, there are likely 3 to 4 missed positive PD cases. Using this optimized combined model and accounting for the estimated prevalence of PD, about ten times the number of individuals would be flagged as high risk or a potential PD case for every real case, indicating that this model is best suited to identifying large groups of individuals to monitor within a health registry or biobank to prioritize for further testing.

The strength of our work is its high balanced accuracy in delineating cases and controls. Its other strength is its applicability and utility across datasets (further details on this in the **Online Appendix** for transfer learning). This model has the potential to be used in large healthcare system settings to identify at-risk individuals for potential monitoring as well as nominating future candidates for various preventative interventions or clinical trial enrollments. Our diagnostic model includes both time-varying (age, UPSIT, and RNAseq data) and time static (biological sex, family history, and genetics) features that likely peak accuracy at the time of diagnosis due to model training on the PPMI dataset, which includes only newly diagnosed imaging-confirmed and unmedicated cases. Additionally, the ability to refine the phenotype of the participant group based on a combination of clinician and algorithmic insight will benefit trial recruitment and could only increase the efficacy of a trial. Finally, since our model is diagnostic in nature and designed to target PD early in the disease course, it may be beneficial in helping get treatments or interventions to patients before irreparable damage has been done as large pools of at risk individuals can be flagged for follow-up and closer monitoring for potential symptom onset ^9^.

We have created an interactive web-based application for others to investigate the driving factors in our best model that incorporates all data modalities; it also gives users the flexibility to explore variations like transcriptomics-only models or a new model with none of the clinico-demographic features present (see **Online Appendix** for details). For the combined model that we have focused on describing in this report; decision plots are provided, these are useful as the web application is capable of letting users explore how and why individuals that were difficult to classify were labeled as cases or controls. The web application (as well as **Figure 4**) shows that the UPSIT score, in general, was the strongest factor in deciding if an individual was classified to be a positive PD case or healthy control by the model. As an example, a decision plot shows that a sample that was clinically diagnosed to be a PD case, we see that most of the features seemed to indicate that the individual was about to be classified as a PD case by the model, but ultimately an unexpectedly high UPSIT score misclassified the individual as a healthy control (decision plots work to visualize the path a model takes before arriving at a classification; see the additional figure in the **Online Appendix** for a graphical representation of misclassification). In general, UPSIT itself accounts for roughly half of the decision making process in our model, and in some instances, is a blessing and a curse with regard to model performance.

Genes and variants affecting the model’s performance shown in **Figure 4** may have some impact on PD biology. Many of the top features we nominated that are shown in **Figure 4** are transcriptomic in nature; this enrichment of transcriptomic data in the top of the feature importance plots may be due to the PRS accounting for a substantial part of the strongest purely genetic aspects of PD risk. Some interesting biologically plausible connections can be drawn from these highly ranked features. For example, the most impactful feature from the transcriptomics data, the expression of gene *HS3ST3A1*, has been implicated in α-Synuclein aggregation in PD cellular models, as well as having been recently part of a novel GWAS finding associated with white matter hyperintensity burden in elderly populations (along with some aspects of cognitive decline) ^10,11^. Another top ranking transcriptomic feature, *OTOL1*, has been suggested before as a putative genetic modifier of familial PD age at onset ^12^. *CHFR* has been associated in previous studies with rotenone related PD risk ^13^. *CASP7* is potentially biologically interesting due to its expression being implicated in apoptosis and neuroprotection as well as rare missense mutations in the gene being associated with late onset familial Alzheimer’s ^14,15^. The genetic variant, rs4238361 is a potential PD risk modifying variant whose nearest coding gene is *VPS13C*, a gene that harbors both rare and common PD variants of interest ^16,17^. The gene *PHF14* has been suggested to be downregulated in neurodegenerative diseases, this potential effect mirrors that suggested in **Figure 4** ^18^. Recent single-cell sequencing work has provided evidence for a connection between *SQLE* and dopamine stress responses in neurons relating to PD risk ^19^. Additionally, *MMP9* overexpression has been suggested to be associated with neuronal cell death in neurodegeneration ^20^.

Another strength of this work is that feature selection from model building easily segues into network community analyses to build relatively low bias networks, compared to those with potential bias taken from literature and text mining ^21,22^. Here we can push therapeutic and biomarker research by identifying communities of connected genes in the blood transcriptome of PD patients. Nodes in these networks suggest shared effects in genetically targeted drugs, informing development cycles and benefitting developers as drugs connected to genetic or genomic data often have a higher level of success in trials compared to those without similar evidence ^23,24^.

Modeling exercises like these not only have the potential to build useful classifiers, they may also identify drug targets. This can happen at the feature selection, and network build phases. In our network community build based on case-only expression data, only two quantitative trait loci in blood from Mendelian randomization in the previously published PD GWAS were included. These two genes are *ZBTB4* and *FCF1*. What may be more interesting is the enriched drug targets within the nominated genes from our analyses of the transcriptomic data. This result is not truly surprising as our network communities are based on genes highly correlated in cases only, and aimed to build clusters of genes connected by similar expression patterns among cases. More interestingly are the findings from the network communities recognizing the overrepresentation of genes targeted by known drugs. For instance, in this study, we uncovered an interaction between gamma hydroxybutyric acid and *SLC16A7, SLC16A3*, and *GABBR1* genes. The *SLC16A7* and *SLC16A3* genes are a part of a family of drug transporter genes known as monocarboxylate transporters^25^. Drug transporters have a role in almost every part of the therapeutic process, from absorption, distribution, and elimination of drug molecules. The *GABBR1* gene encodes a receptor for gamma-aminobutyric acid (GABA) expressed throughout the brain; defects in this gene underlie several neurobehavioral diseases^25,26^. Gamma hydroxybutyric acid acts as an agonist, activativating GABA-B receptors to exert its sedative effects. Identification of drug metabolism and receptor gene/drug interactions may lead to and drug discovery, thereby helping us optimize drug therapy.

Our main weakness in this research is the lack of diversity in available sample series. Current research suggests that genetic predictive models have mixed results when being applied across genetic ancestry groups^27^. With subsequent iterations of this work being facilitated by the Global Parkinson’s Genetics Program (GP2) program over the next five years^28,29^, we hope to expand this modeling effort into a diverse set of genetic ancestry groups and generally in larger sample series. We also acknowledge that no optimal dataset to validate the findings from PPMI exists because of the inherent study design of PPMI focusing on unmedicated recently diagnosed PD cases. Clearly, the ongoing extension of the PPMI study will facilitate further work.

Overall, we believe this work represents a significant conceptual and scientific advance past previous efforts. This classifier has improved performance, is more broadly applicable, and is highly reproducible. Further, the transparency of this approach and the contributions of data types move the field away from black-box predictors of disease. A further strength of this work is the use of open-source automated ML software thoughtfully designed for scientists and scientific data, developing models and validating them on public controlled-access datasets, visualizing the top contributing features, and providing all the code and software publicly.

This work has helped to push past the previous paradigm of focusing on a single biomarker or class of biomarker in biomedical research to maximize data value for clinical and computational scientists by leveraging ML algorithms that explore complex relationships between features. We have provided a model(s) to improve risk prediction in PD to help with interventional and prospective studies as well as healthcare resource prioritization. We have also integrated additional analyses and data resources that may aid in developing and/or refining future interventions.

## CONTRIBUTORS

MBM, FF, and MAN contributed to the concept and design of the study. MBM, HL, DV, HI, IV, EH, DS, JJK, YS, MB, DWC, KVJ, CB, ABS, and MAN were involved in the acquisition of data, data generation, and data cleaning. MBM, HL, DV, HI, FF, AD, LS and MAN did the analysis and interpretation of data. MBM, HL, DV, HI, LS, AD, IV, EH, DS, SBC, JJK, YS, MB, WN, SHH, RHC, MM, JAB, DWC, KVJ, HRM, JAH, CB, ABS, and MAN contributed to the drafting of the article and revising it critically.

## DECLARATION OF INTERESTS

HL, HI, FF, DV, YS, and MAN declare that they are consultants employed by Data Tecnica International, whose participation in this is part of a consulting agreement between the US National Institutes of Health and said company. HRM is employed by UCL. In the last 24 months he reports paid consultancy from Biogen, Biohaven, Lundbeck; lecture fees/honoraria from Wellcome Trust, Movement Disorders Society. Research Grants from Parkinson’s UK, Cure Parkinson’s Trust, PSP Association, CBD Solutions, Drake Foundation, Medical Research Council, Michael J Fox Foundation. HRM is also a co-applicant on a patent application related to C9ORF72 - Method for diagnosing a neurodegenerative disease (PCT/GB2012/052140).

## ACKNOWLEDGEMENTS

This work was supported in part by the Intramural Research Program of the National Institute on Aging and National Institute of Neurological Disorders and Stroke (project number Z01-AG000949-02). Data used in the preparation of this article were obtained from the AMP PD Knowledge Platform. For up-to-date information on the study, https://www.amp-pd.org. AMP PD – a public-private partnership – is managed by the FNIH and funded by Celgene, GSK, the Michael J. Fox Foundation for Parkinson’s Research, the National Institute of Neurological Disorders and Stroke, Pfizer, Sanofi, and Verily. Clinical data and biosamples used in the preparation of this article were obtained from the Parkinson’s Progression Markers Initiative (PPMI), and the Parkinson’s Disease Biomarkers Program (PDBP). PPMI – a public-private partnership – is funded by the Michael J. Fox Foundation for Parkinson’s Research and funding partners, including full names of all of the PPMI funding partners found at www.ppmi-info.org/fundingpartners. The PPMI Investigators have not participated in reviewing the data analysis or content of the manuscript. For up-to-date information on the study, visit www.ppmi-info.org. The Parkinson’s Disease Biomarker Program (PDBP) consortium is supported by the National Institute of Neurological Disorders and Stroke (NINDS) at the National Institutes of Health. A full list of PDBP investigators can be found at https://pdbp.ninds.nih.gov/policy. The PDBP Investigators have not participated in reviewing the data analysis or content of the manuscript. PDBP sample and clinical data collection is supported under grants by NINDS: U01NS082134, U01NS082157, U01NS082151, U01NS082137, U01NS082148, U01NS082133. A portion of the resources used in the preparation of this article were obtained from Global Parkinson’s Genetics Program (GP2). GP2 is funded by the Aligning Science Against Parkinson’s (ASAP) initiative and implemented by The Michael J. Fox Foundation for Parkinson’s Research (https://parkinsonsroadmap.org/gp2/). For a complete list of GP2 members, see https://parkinsonsroadmap.org/gp2/. Workflow diagram was created with BioRender.com.

## Online repository of code and models for transfer learning

### Online Repository

The link to the GitHub repository (https://github.com/GenoML/GenoML_multimodal_PD/) includes the following:

- Figures referenced in the manuscript
- Code used for manuscript
- Tables referenced in the manuscript
- Additional supplementary figures and tables
- Links and references to the main software used (GenoML)
- Trained and tuned models and their associated performance metrics for each model referenced in the manuscript for the community to be able to deploy and use
- An example of how to run these trained and tuned models for transfer learning
- Link to the interactive web application for the community to investigate further

### Network Communities

The network community code can be found on the **Online Repository** here. Annotated community members plus additional sparse annotations from Nalls et al. 2019 can be found here. Page ranks and similar annotations for genes comprising the network communities can be found here.

## Online Methods

### Study design and participants

Clinical, demographic plus genome-wide DNA and RNA sequencing data were taken at baseline visits from the Parkinson’s Progression Marker Initiative (PPMI) and the Parkinson’s Disease Biomarkers Program (PDBP) in cases with PD and controls unaffected by neurologic diseases. Since our model is retrospective, we aimed only to analyze refined Parkinson’s disease diagnosis, by excluding any samples with conflicting diagnostic data within a decade of post-enrollment follow-up. We excluded any case whose medical history included an additional neurological disease diagnosis or retraction of their PD diagnosis during follow-up. We also excluded controls developing PD or another neurodegenerative disease(s) after enrollment. Additionally, a subset of Parkinson’s disease cases and controls from the PPMI study were excluded as they came from a targeted study recruitment design purposely enriching for known genetic risk mutation carriers (*LRRK2* and *GBA* mutation carrier focused recruitment).

Participants with required clinical, demographic and genomic (DNA and RNA sequencing) data were identified for inclusion, with excessive missing data (> 15% per feature) as exclusion criteria. Each contributing study abided by the ethics guidelines set out by their institutional review boards, and all participants gave informed consent for inclusion in both their initial cohorts and subsequent studies.

Clinical and demographic data ascertained as part of this project included age at diagnosis for cases and age at baseline visit for controls. Family history (self-reporting if a first or second-degree relative has a Parkinson’s disease diagnosis) was also a known clinico-demographic feature of interest. Ashkenazi Jewish status was inferred using principal component analysis comparing those samples to a genetic reference series ^1^. Sex was clinically ascertained but also confirmed using X chromosome heterozygosity rates. The University of Pennsylvania Smell Inventory Test (UPSIT) was used in modeling ^2^. For a summary of basic clinical and demographic features, please refer to **Table 1**.

DNA sequencing data were generated using Illumina’s standard short-read technology, and the functional equivalence pipeline during alignment was the Broad Institute’s implementation^3^. Jointly genotyped sequencing data using the standard GATK pipeline from AMP-PD was used. This process is described in detail, from sample prep to variant calling, in a separate manuscript detailing the AMP PD whole-genome DNA sequencing effort [under review] ^4^.

Quality control for these samples based on genetic data output by the pipeline included the following inclusion criteria: concordance between genetic and clinically ascertained genders, call rate > 95% at both the sample and variant levels, heterozygosity rate < 15%, freemix estimated contamination rate < 3%, transition:transversion ratio > 2, unrelated to any other sample at a level of the first cousin or closer (identity by descent < 12.5%) and genetically ascertained European ancestry. For inclusion of whole-genome DNA sequencing data, the variants must have passed basic quality control as part of the initial sequencing effort (PASS flag from the joint genotyping pipeline) as well as meeting the following criteria: non-palindromic alleles, missingness by case-control status P > 1E-4, missingness by haplotype P > 1E-4, Hardy-Weinberg p-value > 1E-4, minor allele frequency in cases > 5% (in the latest Parkinson’s disease meta-GWAS) ^5^. As an *a priori* genetic feature to be included in our modeling efforts, we also used the basic polygenic risk score from the latest Parkinson’s disease meta-GWAS (genome-wide significant loci only) that did not include our testing or training samples as weights ^5^.

RNA sequencing data from whole blood on the same samples was generated by the Translational Genomics Research Institute team using standard protocols for the Illumina NovaSeq technology ^6^. For this study, we focused on blood withdrawn at baseline. Variance stabilized counts were adjusted for experimental covariates using standard limma pipelines^7^. Gene expression counts for protein-coding genes were extracted, then differential expression p-values were calculated between cases and controls using logistic regression adjusted for additional covariates of sex, plate, age, ten principal components, and percentage usable bases.

This project is carried out entirely within an open science framework, and the code and data underlying the summary above can be found in linked notebooks and datasets described in the **Online Repository**.

### Data munging

As part of the initial data munging, principal components summarizing genetic variation in DNA and RNA sequencing data modalities are generated separately. For the DNA sequencing, ten principal components were calculated based on a random set of 10,000 variants sampled after linkage disequilibrium pruning that kept only variants with r2 < 0.1 with any other variants in +/− 1MB. As a note, these variants were not p-value filtered based on recent GWAS, but they do exclude regions containing large tracts of linkage disequilibrium^8^. For RNA sequencing data, all protein-coding genes’ read counts per sample were used to generate a second set of 10 principal components. All potential features representing genetic variants (in the form of minor allele dosages) from sequencing were then adjusted for the DNA sequence-derived principal components using linear regression, extracting the residual variance. The same was done for RNA sequencing data using RNA sequencing derived principal components. This way, we statistically account for latent population substructure and experimental covariates at the feature level to increase generalizability across heterogeneous datasets. In its simplest terms, all transcriptomic data were corrected for possible confounders, and the same is done for genotype dosages. After adjustment, all continuous features were then Z transformed to have a mean of 0 and a standard deviation of 1 to keep all features on the same numeric scale when possible. Once feature adjustment and normalization were complete, internal feature selection was carried out in the PPMI training dataset using extra decision trees to identify features contributing information content to the model while reducing the potential for overfitting ^9,10^.

### Feature and model selection

After the data munging process (quality control, feature selection, adjustment, and scaling) described above, data from PPMI was randomly split into 70% training and 30% testing. Training of the algorithms was performed on the training set, and validation of the algorithms was performed on the testing set. A total of 12 well-performing ML algorithms were competed to identify which algorithm could maximize AUC across the two classes (cases and controls). These algorithms were chosen due to their success in other domains, execution in Python’s scikit-learn package, and their ability to export probability-based predictions, allowing the training, testing, and interpretation of the model more straightforward. The algorithms included are: logistic regression (LogisticRegression), random forests (RandomForestClassifier), adaptive boosting (AdaBoostClassifier), gradient boosting (GradientBoostingClassifier), stochastic gradient descent (SGDClassifier), support vector machines (SVC), multi-layer perceptron neural networks (MLPClassifier), k-nearest neighbors (KNeighborsClassifier), linear discriminant analysis (LinearDiscriminantAnalysis), quadratic discriminant analysis (QuadraticDiscriminantAnalysis), bagging (BaggingClassifier) and extreme gradient boosting (XGBClassifier). The algorithm with the highest AUC and balanced accuracy in the withheld 30% of PPMI was selected for tuning and cross-validation. Tuning is the process in which multiple algorithm hyperparameters, such as learning rate, are tested to optimize performance. The best hyperparameters were chosen through cross-validation, a technique that estimates model performance on unseen data by training and testing the model on different splits of the dataset. The top competing algorithm was then selected to undergo a computationally intensive hyperparameter tuning phase in the entire PPMI dataset, no longer split into training and testing once, instead, undergoing cross-validation each time parameters were iterated. In this analysis, the top-performing algorithm (AdaBoostClassifier) was tuned for several potential predictors (estimators) between 1 and 1000 for 25 random iterations at 5-fold cross-validation per iteration.

This process detailed in the paragraphs above was carried out 49 times, at varying thresholds of p-values based feature inclusion thresholds. We iterated across all possible combinations of p-value thresholds [1E-2, 1E-3, 1E-4, 1E-5, 1E-6, 1E-7, 1E-8] for genetic data from the most recent published GWAS and for transcriptomic data from our differential expression work also described above^5^. Once the pipeline above was run for all 49 threshold combinations, we picked the p-value thresholds and algorithm that performed best in the training dataset for tung at cross-validation and external validation in PDBP to evaluate its generalizability and performance (all 49 models are available in the **Online Repository**). Prior to starting the modeling process, we specified that this manuscript would focus on the model trained in PPMI that presented the highest mean AUC, accuracy, and balanced accuracy in withheld samples before moving on to validation in a *de novo* dataset. We acknowledge that incorporating p-values as a pre-filtering step in the feature selection phase may cause data leakage to some degree. The PPMI training set is only a minuscule portion of the most recent GWAS study (less than 0.1% of the sample size); additionally, the algorithmic feature selection described above is generally much more conservative and excluded the majority of features reaching the p-value thresholds of interest. In this report, we only focused on a model with a 1E-5 maximum p-value for genetic data inclusion and a 1E-2 maximum P for transcriptomic data inclusion; however, all potential models were exported and saved for public use in transfer learning for similar datasets for the scientific community (**Online Repository**).

### Post-hoc optimization for class imbalance

After training, we refit the model to the withheld samples using an optimized threshold for case probability based on Youden’s J calculation to better account for case-control imbalance and subsequently increase balanced accuracy and related metrics^11^. This *post-hoc* optimization was done again after fitting the tuned model to the external validation cohort.

### Network communities

After building the ML classifiers, we turned our attention to potentially novel PD gene networks that may be hidden within the classifier’s selected features. First, we extracted all identified RNA feature counts for this subset of genes. Next, we subsetted to cases only and calculated the correlation between gene-level transcriptomic data for the nominated genes to build a graph space (minimum correlation coefficient (r), the threshold of 0.8 for connections between gene nodes). Then the Leiden algorithm was implemented to cluster the genes within the larger network into related communities; finally, we calculated a modularity score to evaluate the quality of our network clusters ^12^.

Looking for potential therapeutic connections across the communities within our defined networks, we utilized webGestaltR; we used its over-representation analysis function to explore druggable target enrichments for network genes within the two available drug databases hosted on the website (DrugBank and GLAD4U) ^13,14,15^. These queries were made under default settings. First, we queried the 300 genes comprising our network communities against a background of all 598 genes nominated at the initial feature selection phase in the transcriptomics data. This estimates how genes comprising our network communities, which are highly correlated in cases, might be enriched compared to genes potentially related to case:control differences. We also looked for enrichments similarly comparing all 598 potential genes delineating cases and controls to > 18,000 protein-coding genes. This was then repeated for our 300 network community genes, investigating over-representation of druggable targets against all protein-coding genes. Our primary goal with this analysis was to see if any drug-related annotations were enriched in our network communities based on correlated gene expression between cases compared to other protein-coding genes that were selected as potential case:control classifying features. This database was accessed on February 23rd, 2021.

### ML metrics and model interpretation

Our prioritized metric for evaluating model performance was the area under the curve, AUC. The AUC metric is an aggregate metric summarizing the performance of a classifier across all potential probability thresholds for delineating cases:controls (the labels used in this study) and is less affected by class imbalance than other common metrics. In general, an AUC of greater than 80% may be considered qualitatively to be a robust and “very good to excellent” diagnostic^16^. For other metrics such as sensitivity and true positive rate (these relate to the proportion of true positive cases identified) or specificity and true negative rate (these relate to the proportion of true controls identified), their values are easily altered by a change to the probability threshold used to split cases and controls. For example, after the model outputs a probability estimate of a sample being a case, a researcher has the option to use the default probability threshold of 50% for binary classification, or use several methods to optimize this threshold for better performance. As part of our automated ML workflow, we output performance metrics at default, followed by optimized probability thresholds (using Youden’s J). We prioritized secondary performance metrics of interest in addition to AUC; these were accuracy and balanced accuracy, the former being the rate of correct predictions, the latter being the mean accuracy weighted across cases, and controls samples used for dealing with imbalanced datasets ^17^.

The Shapley additive explanations (SHAP) approach was used to evaluate each feature’s influence in the ML model. This approach, used in game theory, assigns an importance (Shapley) value to each feature to determine a player’s contribution to success ^18^. In this analysis, the concept of “success” has been replaced by case status. Shapley explanations enhance understanding by creating accurate explanations for each observation in a dataset. The SHAP package was used to calculate and visualize these Shapley values seen in the figures in the manuscript and the interactive website ^19,20^. A surrogate xgboost model was trained in 70% of the data, and later tested in the 30% of withheld data to evaluate the model’s contributing features. The interactive website (https://share.streamlit.io/anant-dadu/shapleypdpredictiongenetics/main) was developed as an open-access and cloud-based platform for researchers to investigate the top features of the model developed in this study and how these may influence the classification (or in some cases, misclassification) of a particular sample. In its simplest description, the Shapley values are similar to standard regression derived relative importance measures with regard to interpretation, and all the Shapley values sum to a total value of 1.

## Details on data from the Accelerating Medicines Partnership - Parkinson’s Disease (AMP-PD) and the Global Parkinson’s Genetic Project (GP2)

### The summaries below are from AMP PD’s website on the different studies used in this manuscript

#### Cohort Summaries - PPMI

##### Study Overview for PPMI

The Parkinson’s Progression Markers Initiative (PPMI) is a study sponsored by the Michael J. Fox Foundation. It is a longitudinal, observational study where participants can contribute clinical, demographic, and imaging data alongside biological samples used for whole-genome sequencing, whole blood RNA sequencing, and other assays at 33 clinical sites globally. PPMI follows participants for anywhere from five to 13 years. For this manuscript, we have only focused on data collected at baseline. This data is now hosted as part of AMP PD’s version 1 release. For more information on the PPMI study, please visit this link (https://amp-pd.org/unified-cohorts/ppmi#study-overview).

##### Study Inclusion Criteria for PPMI - PD Cases

1. Patients must have at least two of the following: resting tremor, bradykinesia, rigidity (must have either resting tremor or bradykinesia); OR either asymmetric resting tremor or asymmetric bradykinesia
2. A diagnosis of Parkinson disease for 2 years or less at Screening
3. Hoehn and Yahr stage I or II at Baseline
4. Confirmation from imaging core that screening dopamine transporter SPECT scan is consistent with dopamine transporter deficit (or for sites where DaTSCANTM is not available, that VMAT-2 PET scan is consistent with VMAT deficit)
5. Not expected to require PD medication within at least 6 months from Baseline. Male or female age 30 years or older at time of PD diagnosis

##### Study Exclusion Criteria for PPMI - PD Cases

1. Currently taking levodopa, dopamine agonists, MAO-B inhibitors, amantadine, or other PD medication
2. Has taken levodopa, dopamine agonists, MAO-B inhibitors, or amantadine within 60 days of Baseline
3. Has taken levodopa or dopamine agonists prior to Baseline for more than a total of 60 days
4. Received any of the following drugs that might interfere with dopamine transporter SPECT imaging: Neuroleptics, metoclopramide, alpha methyldopa, methylphenidate, reserpine, or amphetamine derivative, within 6 months of Screening
5. Current treatment with anticoagulants (e.g., coumadin, heparin) that might preclude safe completion of the lumbar puncture
6. Condition that precludes the safe performance of routine lumbar puncture, such as prohibitive lumbar spinal disease, bleeding diathesis, or clinically significant coagulopathy or thrombocytopenia
7. Use of investigational drugs or devices within 60 days prior to Baseline (dietary supplements taken outside of a clinical trial are not exclusionary, e.g., coenzyme Q10)

##### Study Inclusion Criteria for PPMI - Healthy Controls

Healthy controls for the PPMI study included males or females 30 years or older at Screening

##### Study Exclusion Criteria for PPMI - Healthy Controls

1. Current or active clinically significant neurological disorder (in the opinion of the Investigator).
2. First degree relative with idiopathic PD (parent, sibling, child)
3. MoCA score < 26
4. Received any of the following drugs that might interfere with dopamine transporter SPECT imaging: Neuroleptics, metoclopramide, alpha methyldopa, methylphenidate, reserpine, or amphetamine derivative, within 6 months of Screening
5. Current treatment with anticoagulants (e.g. coumadin, heparin) that might preclude safe completion of the lumbar puncture
6. Condition that precludes the safe performance of routine lumbar puncture, such as prohibitive lumbar spinal disease, bleeding diathesis, or clinically significant coagulopathy or thrombocytopenia
7. Use of investigational drugs or devices within 60 days prior to baseline (dietary supplements taken outside of a clinical trial are not exclusionary, e.g., coenzyme Q10)

#### Cohort Summaries - PDBP

##### Study Overview for PDBP

The Parkinson’s Disease Biomarkers Program (PDBP) is a study sponsored by the National Institute of Neurological Disorders and Stroke (NINDS). It is a longitudinal, observational study where participants can contribute clinical, demographic, and imaging data alongside biological samples used for whole-genome sequencing, whole blood RNA sequencing, and other assays. The goal of this study is to accelerate the discovery of promising new diagnostic and progression biomarkers for Parkinson’s Disease. This data is now hosted as part of AMP PD’s version 1 release. For more information on the PDBP study, please visit this link (https://amp-pd.org/unified-cohorts/pdbp#study-overview).

##### Study Inclusion Criteria for PDBP - PD Cases

1. Clinically diagnosed with Parkinson’s Disease
2. Male or Female aged 21 years or older at screening
3. Able to cooperate with consent procedures (or has appropriate surrogate as defined and approved per local IRB)
4. Able to participate in study activities including all required clinical assessments and biological donations
5. Participation would not lead to hardship or adverse health or mental health conditions

##### Study Exclusion Criteria for PDBP - PD Cases

1. Clinical Diagnosis uncertain at the time of enrollment
2. Condition that preclude the safe performance of routine lumbar puncture, such as prohibitive lumbar spinal disease, bleeding diathesis, or coagulopathy or thrombocytopenia
3. Current treatment with anti-coagulants (e.g., Coumadin, heparin) that might preclude safe completion of the lumbar puncture
4. Has a history of neuroleptic use or exposure
5. Has a history of schizophrenia
6. Otherwise unable to participate in biological specimen collection due to a medical condition or medication status (other than items listed above)
7. Otherwise unable to participate in clinical assessments due to a medical condition or medication status (other than items listed above)
8. Unable to participate in consent procedures
9. Use of investigational drugs or devices within 60 days prior to baseline visit (dietary supplements such as Coenzyme Q10, for example, are not exclusionary)

##### Study Inclusion Criteria for PDBP - Healthy Controls

1. Male or Female aged 21 years or older at screening
2. Able to cooperate with consent procedures (or has appropriate surrogate as defined and approved per local IRB)
3. Able to participate in study activities including all required clinical assessments and biological donations
4. Participation would not lead to hardship or adverse health or mental health conditions

##### Study Exclusion Criteria for PDBP - Healthy Controls

1. Has a current or clinically significant neurological disorder in the opinion of the investigator
2. Family history of Neurodegenerative disease in a first degree relative or second degree blood relative
3. Condition that preclude the safe performance of routine lumbar puncture, such as prohibitive lumbar spinal disease, bleeding diathesis, or coagulopathy or thrombocytopenia
4. Current treatment with anti-coagulants (e.g., Coumadin, heparin) that might preclude safe completion of the lumbar puncture
5. Has a history of neuroleptic use or exposure
6. Has a history of schizophrenia
7. Otherwise unable to participate in biological specimen collection due to a medical condition or medication status (other than items listed above)
8. Otherwise unable to participate in clinical assessments due to a medical condition or medication status (other than items listed above)
9. Unable to participate in consent procedures
10. Use of investigational drugs or devices within 60 days prior to baseline visit (dietary supplements such as Coenzyme Q10, for example, are not exclusionary)

### Age distribution per cohort

**Supplementary Figure 1:**
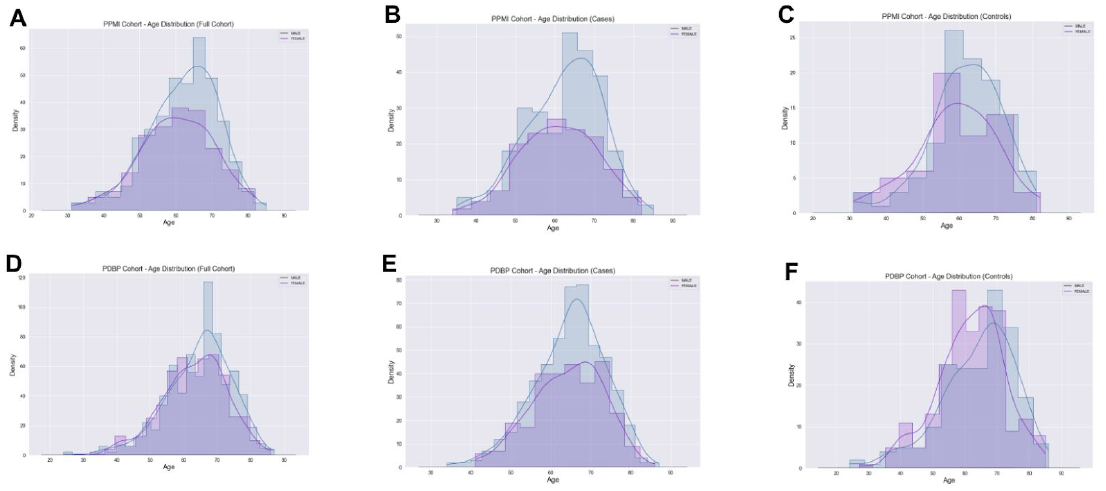
Age distributions of cohorts, broken down by cases and controls | Blue indicates male, while purple indicates female. Panels A, B, and C show the age distribution of males and females in the full PPMI cohort, cases in PPMI, and controls in PPMI, respectively. Panels D, E, and F show the age distribution of males and females in the full PDBP cohort, cases in PDBP, and controls in PDBP, respectively.

### Network Graphical Summary of Nominated Genes

We identified 13 network communities consisting of 300 genes with an Erdos-Renyi modularity score of 0.794 (a modularity score closer to 1 indicates better model fit).

**Supplementary Figure 2:**
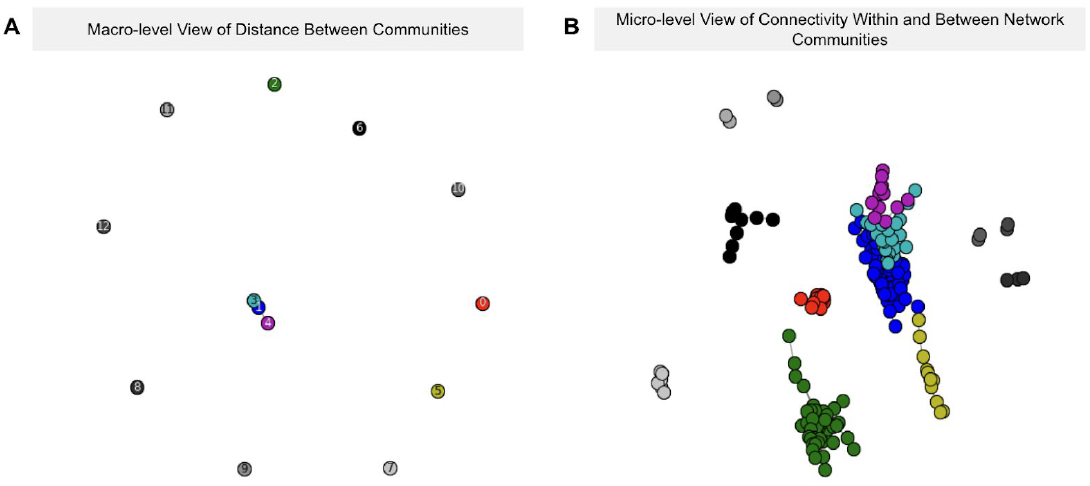
Network plot of nominated genes | Panel A provides a macro-level view of the distance between communities (color-coded). Panel B is a micro-level view of connectivity within and between network community modules. The colors of communities in Panel A correspond to those in panel B.

## Misclassified cases by the best performing model

### Encrypted ID of Misclassified Case: 756ac1345d7068cdc60c8b2583a80092

Decision plots work on visualizing the path a model takes before arriving at a classification. A decision plot shows that a sample that was clinically diagnosed to be a PD case, we see that most of the features seemed to indicate that the individual was about to be classified as a PD case by the model, but ultimately an unexpectedly high UPSIT score misclassified the individual as healthy control.

**Supplementary Figure 3:**
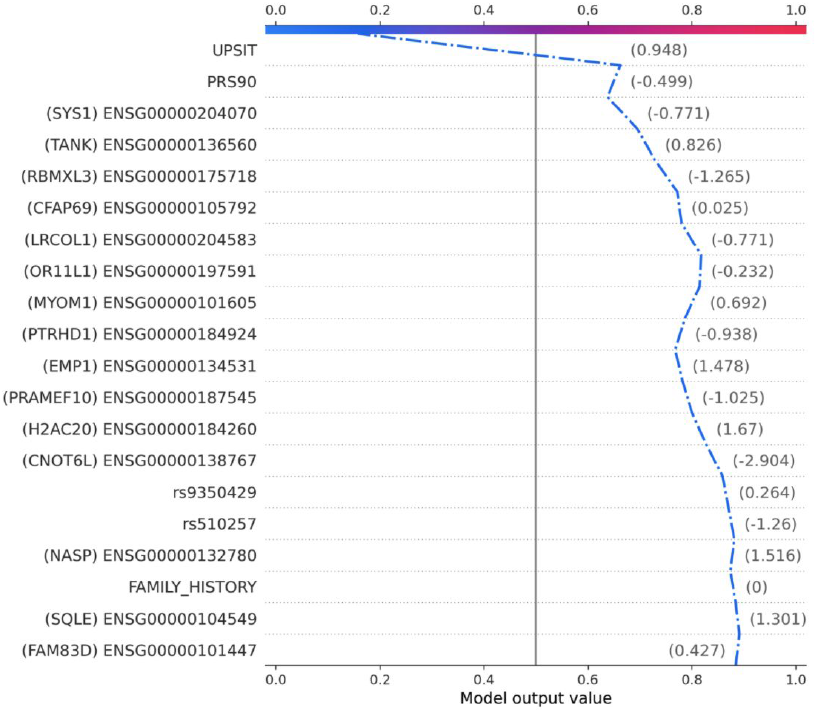
Misclassified case as a healthy control using the best model

## Notes

https://github.com/GenoML/GenoML_multimodal_PD

https://amp-pd.org/

https://genoml.com/

https://share.streamlit.io/anant-dadu/shapleypdpredictiongenetics/main

